# Genetic estimates and genome-wide association studies of antibody response in Tanzanian dairy cattle

**DOI:** 10.1101/2024.08.05.606566

**Authors:** Luis E Hernandez-Castro, Elizabeth Anne Jessie Cook, Oswald Matika, Isaac Joseph Mengele, Shabani Kiyabo Motto, Shedrack Festo Bwatota, Bibiana Zirra-Shallangwa, Ricardo Pong-Wong, James Prendergast, Raphael Mrode, Philip G. Toye, Daniel Mushumbusi Komwihangilo, Eliamoni Lyatuu, Benedict E. Karani, Getrude Nangekhe, Okeyo Ally Mwai, Gabriel Mkilema Shirima, Barend Mark de Clare Bronsvoort

**Affiliations:** Centre for Tropical Livestock Genetics and Health (CTLGH), Roslin Institute, University of Edinburgh, Easter Bush Campus, EH25 9RG, United Kingdom; The Roslin Institute and The Royal (Dick) School of Veterinary Studies, The University of Edinburgh, Easter Bush Campus, Midlothian, EH25 9RG, United Kingdom; International Livestock Research Institute (ILRI), Nairobi, Kenya; Centre for Tropical Livestock Genetics and Health (CTLGH), ILRI Kenya, P.O. Box 30709, Nairobi 00100, Kenya; Department of Global Health and Bio-Medical Sciences, School of Life Science and Bioengineering, The Nelson Mandela African Institution of Science and Technology, Arusha, Tanzania; Tanzania Veterinary Laboratory Agency, Central Veterinary Laboratory, Dar es Salaam, Tanzania; Scotland’s Rural College, Easter Bush Campus, Roslin Institute, Midlothian, EH25 9RG, United Kingdom; Tanzania Livestock Research Institute (TALIRI), Dodoma, Tanzania; International Livestock Research Institute (ILRI), Dar es Salaam, Tanzania

**Keywords:** Genetic estimates, GWAS, genomic population characterisation, Tanzanian dairy cattle

## Abstract

Identifying the genetic determinants of host defence against infectious pathogens is central to enhancing disease resilience and therapeutic efficacy in livestock. Here we have taken a genome-wide association approach to identify genetic variants associated with the presence of serological markers for important infectious diseases affecting dairy cattle in smallholder farms. Assessing 668,911 single-nucleotide polymorphisms in 1977 crossbreed cattle sampled from six regions of Tanzania, we identified high levels of interregional admixture and European introgression which may increase infectious disease susceptibility relative to indigenous breeds. Heritability estimates ranged from 0.03 (SE ± 0.06) to 0.44 (SE ± 0.07) depending on the pathogen assayed. Preliminary genome scans revealed several loci associated with seropositivity to the viral diseases Rift Valley fever and bovine viral diarrhoea, the protozoan parasites *Neospora caninum* and *Toxoplasma gondii*, and the bacterial pathogens *Brucella sp, Leptospira hardjo* and *Coxiella burnetti*. The associated loci mapped to genes involved in immune defence, tumour suppression, neurological processes, and cell exocytosis. We discuss future work to clarify the cellular pathways contributing to general and taxon-specific infection responses and to advance selective breeding and therapeutic target designs.

## Introduction

The smallholder dairy cattle sector in Tanzania contributes greatly to the country’s gross domestic product, as well as in securing Tanzanian households’ food, income and employment [^1–3^]. Despite control efforts, animal disease remains a significant constraint to productivity and profitability in the sector with estimated mortality of up to 14% across herds [^4,5^]. Cattle pathogens causing abortion (e.g., Rift Valley Fever virus (RVFV), Bovine viral diarrhea virus (BVDV) and *Neospora caninum*) and/or zoonosis (e.g., *Brucella abortus*, *Leptospira* hardjo, *Coxiella Burnetii* and *Toxoplasma gondii*) contribute greatly to economic loss, threaten human public health and are currently circulating at considerable levels across Tanzania regions [^6–9^].

The defence processes (resilience) by which organisms limit pathogen loads (resistance) or the damage caused by given pathogen loads (tolerance) are crucial to the epidemiology of infectious diseases [^10–12^]. Changes in host immunological and pathogenesis mechanisms can particularly reduce transmission, and therefore prevalence, by blocking infection or eliminating pathogens [^13,14^]. Natural host genetic variability (e.g., α-thalassaemia, sickle cell haemoglobin and G6PDH deficiency) has been shown to protect against deadly *Plasmodium falciparum* infection [^15–18^]. Several genetic variants and genes (e.g., HLA and IFNAR2 genes) linked to host pathophysiological processes have been associated to SARS-CoV-2 susceptibility and COVID-19 severity [^19,20^]. Recently, a CRISPR/Cas9-mediated knockout to alter the BVDV binding domain of the CD46 gene evidenced reduced susceptibility in a cloned calf [^21,22^]. Therefore, understanding the genetic bases of disease resilience could complement current control strategies and reduce the burden of endemic or emerging infectious diseases [^12,23^].

Given limited resources and feasibility to deploy disease control options (e.g., vaccination, biosecurity, contact tracing, etc), health improvement of smallholder dairy cattle via genomic selection could complement conventional disease control efforts in Tanzania and other low and middle-income countries (LMICs). Host antibody response due to recent infection or previous exposure has been shown to be heritable, and the genetic factors influencing these traits have been explored for several human infectious pathogens [^24–26^]. Heritability of antibody response to infectious diseases has also been described in several livestock species. For example, Liu *et al.* (2014) [^27^] reported heritability estimates of 0.36 (± 0.075) and 0.35 (± 0.077) respectively for antibody response to Newcastle disease and avian influenza virus in poultry. In a population of Danish dairy cattle, heritability estimates of antibody response to *Mycobacterium avium* subsp. *paratuberculosis*, which causes diarrhoea and decreases milk yield, was estimated to be 0.10 (± 0.05) [^28^]. Moderate heritability estimates of 0.32 (± 0.09) for immune competence after vaccination against *Clostridium tetani* were reported in Angus beef calves suggesting an opportunity for immune competency improvement via genetic selection [^29^]. Antibody response to bovine herpesvirus-1 (BoHV-1), which causes latent infectious bovine rhinotracheitis (IBR), was heritable in Irish cattle herds with values ranging from 0.12 (± 0.05) to 0.14 (± 0.04) [^30^]. Including health-related traits (e.g., humoral immune response) to breeding programmes could assist in reducing disease burden in livestock.

Any quantitative trait locus (QTL) regions identified as associated with disease resistance could guide breeding, genome editing or diagnostics and therapeutic programmes. Resistance to *Mycobacterium bovis* has been attributed to several regions in the *Bos taurus* genome such as loci containing the PTPRT and MYO3B genes [^31^]. Tolerance to the protozoan parasite *Theileria parva* (East Coast fever -ECF) in cattle has been linked to a locus spanning a paralogue of the FAF1 gene, which significantly increased the survival odds in animals with a tolerance allele [^32^]. Identification of candidate genes has allowed editing bovine genomes to express resistance to pathogens such as *M. bovis* or BVDV [^33^].

The aim of our current study was to investigate the antibody response variation against zoonotic and reproductive cattle infectious diseases, as well as their genetic parameter estimates, and a preliminary genome-wide association study to identify potential QTL regions of importance in health improvement breeding programmes. We carried out population structure, heritability estimates, and genome-wide association studies with 668,911 SNPs in 1977 crossbred smallholder dairy cattle from six regions in Tanzania.

## Results

### Tanzanian small holder cattle, genotyping, and serology traits

A total of 2045 crossbred cattle were sampled across six regions important for dairy production in Tanzania (Fig 1A) and tested for antibodies to seven pathogens. Animals were first genotyped with a mid-density (100K) array, and then, the genotypes were imputed to high-density (∼ 600K) (see Methods). Thirty-four animals were removed as they failed the quality control (QC) measures from the imputation pipeline due to high levels of missing SNPs, and a further 34 were removed due to high levels of parent-offspring relatedness. The call rate (> 90%) and minor allele frequency (> 0.01) thresholds removed 17,132 of the imputed SNPs. Following imputation and QC measures of the initial 99,229 SNPs, 668,911 remained for subsequent analyses. In the final 1,977 cattle dataset, 25.2% of the animals were seropositive to bovine viral diarrhoea virus (BVDV), 19.1% to *Neospora caninum*, 13.2% to *Leptospira* hardjo, 9.3% Rift Valley Fever virus (RVFV), 5.9% to *Toxoplasma gondii*, 3.9% to *Coxiella Burnetii*, and 2.4% to *Brucella abortus* (Fig 1B).

**Fig 1.**
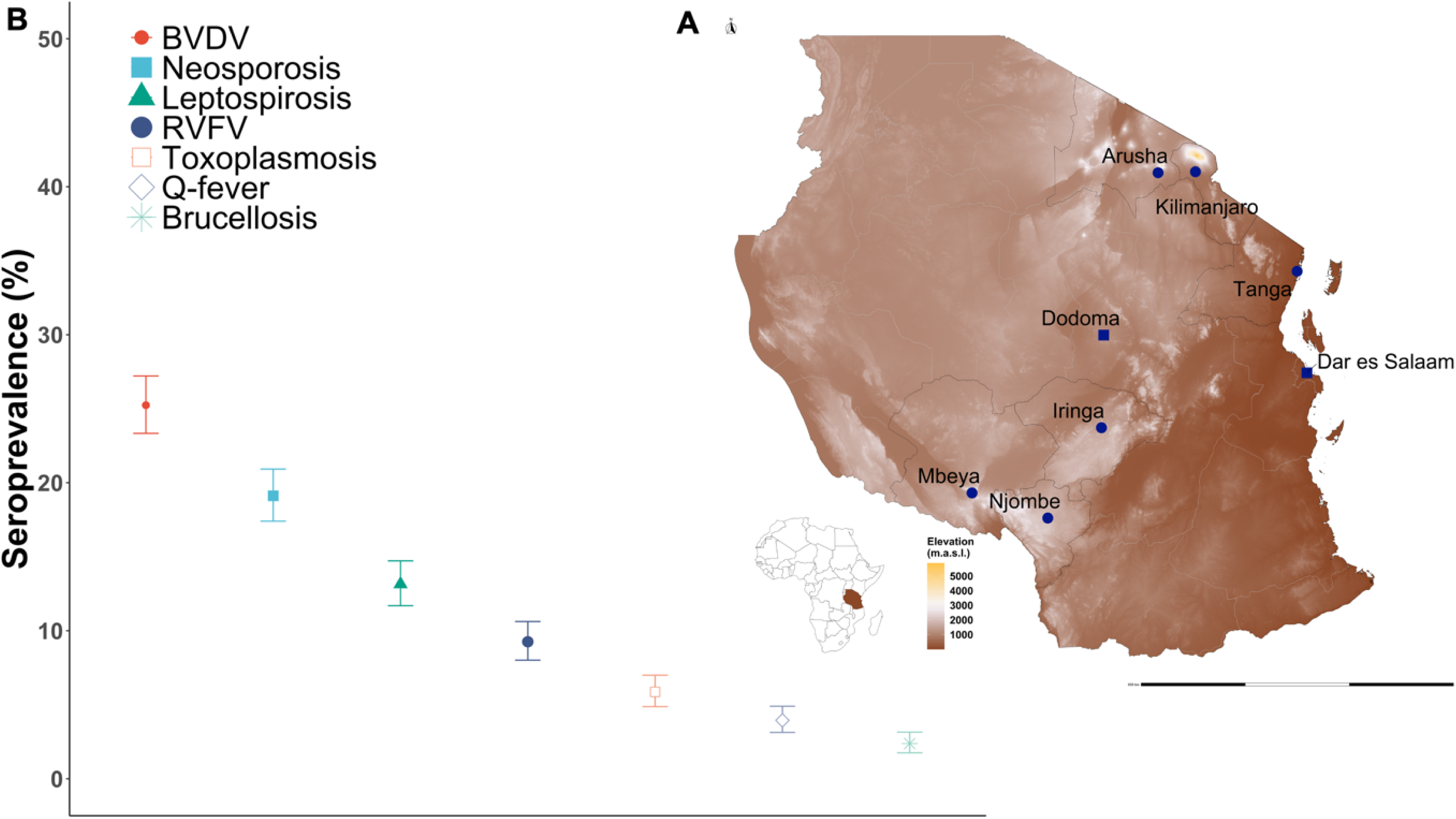
Seroprevalence of reproductive and zoonotic diseases among smallholder dairy cattle in Tanzania. **A**, geographic location of the six regions sampled in Tanzania: Arusha, Kilimanjaro, Tanga, Iringa, Mbeya, and Njombe. **B**, seroprevalence to seven pathogens among smallholder dairy cattle is variable with the highest seropositivity to BVDV, Neosporosis and Leptospirosis. Source map: www.usgs.gov/centers/eros/science/usgs-eros-archive-digital-elevation-global-multi-resolution-terrain-elevation.

### Population structure and ancestry

To investigate the population stratification and structure among these crossbreed Tanzanian cattle (TZA) and their genomic context with a reference of closely pure breeds we performed principal components analysis (PCA), and ADMIXTURE analysis. The reference cattle populations included closely European (ET; Holstein and Jersey, n=99) and African taurine (AT; N’Dama and Muturu, n= 59), and Asian zebu (AZ; Gir and Nelore, n= 65) populations [^32^] (Fig 2A).

**Fig 2.**
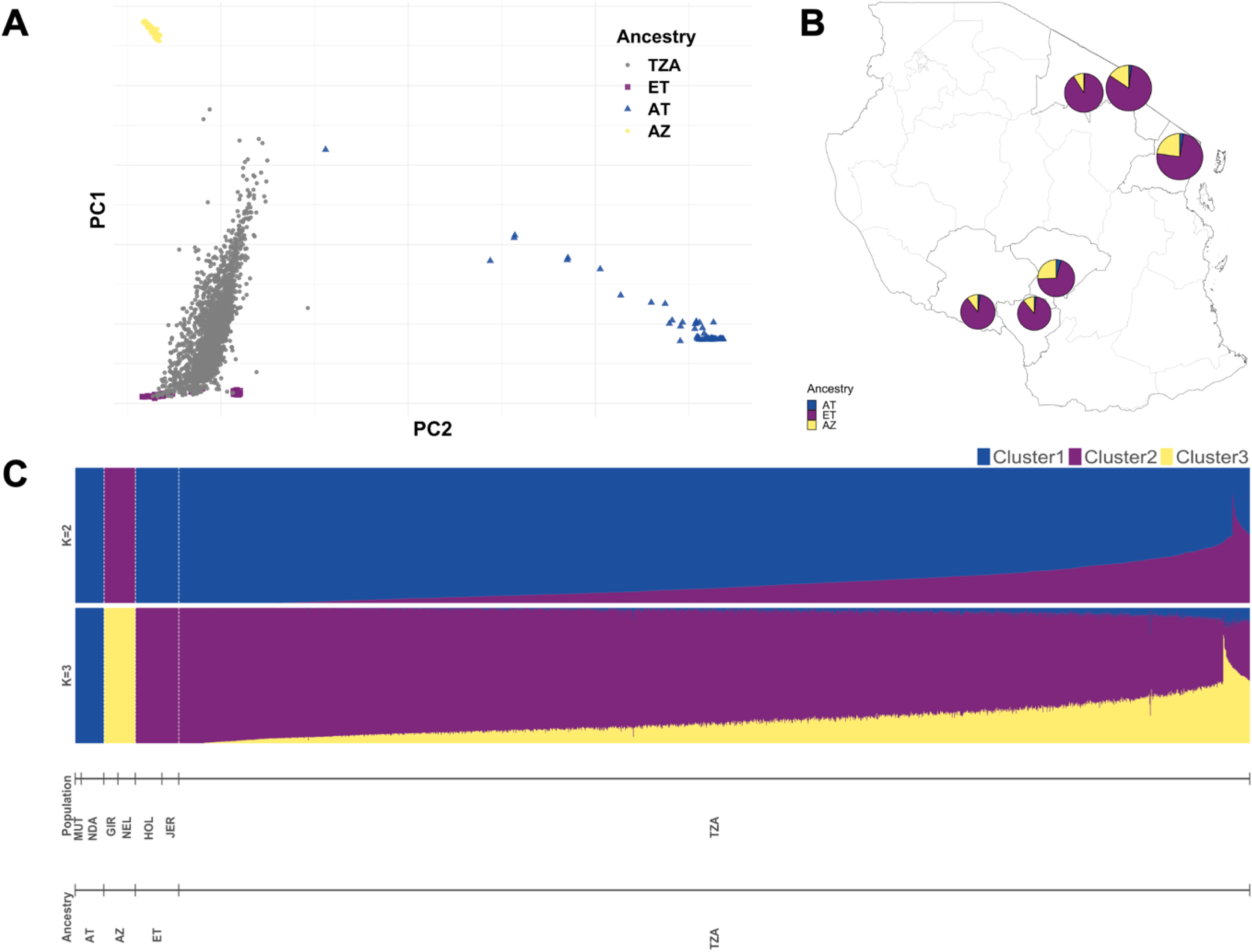
Population structure and ancestry among smallholder dairy cattle in Tanzania. **A**, the scatter plot shows Tanzanian cattle samples (grey dots) within reference populations (European taurine (ET): Holstein and Jersey; African taurine (AT): N’Dama and Muturu; Asian zebu (AZ): Gir and Nelore). **B**, Pie charts show the percentage ET, AT, and AZ ancestry among Tanzanian cattle across the six sampled regions estimated using a supervised admixture analysis, **C**, which indicated high levels of European introgression across samples.

First, we visualised the first two principal components (PCs) of a principal component analysis (PCA) with a reference set of pure breeds. The first and second PCs explained 47.2% and 10.9% of the total genomic variation respectively (S1 Fig). Roughly five clusters were defined in the PCA scatter plot, in which Tanzanian samples grouped across European taurine (ET; Holstein and Jersey samples were closely separated in two) and Asian zebu (AZ; Gir and Nelore clustered together) samples. African taurine samples (AT; N’Dama and Muturu clustered towards PC2 axis) clustered separately from the rest of the populations.

Supervised admixture analysis estimated the proportion of predefined ancestral populations in our Tanzanian samples (Fig 2C). At *K* = 2 predefined ancestral populations, *Bos taurus* and *Bos indicus* ancestry is evident at variable levels across the Tanzanian cattle. The proportion of African and European taurine ancestry diverged in Tanzanian samples when assuming *K* = 3 fixed ancestral populations. Across all sampled regions in Tanzania, animals had high levels of European ancestry, followed by Asian zebu and least African taurine (Fig 2B). Unsupervised admixture analysis revealed similar patterns of ancestries at *K* = 2 and *K* = 3 (S2 Fig), and more than seven clusters (accounting for six reference population and our Tanzanian population) were likely present in our data set as suggested in the cross-validation plot (S3 Fig).

### Heritability estimates of serological responses

We estimated the heritability of serological response to seven pathogens commonly causing abortion in cattle using linear mixed models (see Methods). The seroprevalences in the cattle population under study was low ranging from 2% for *Brucella abortus* to 25% for BVDV. The heritability estimates assuming a binary trait using the observed scale (h^2^_o_) ranged from 0.03 (SE ± 0.06) for Q-fever to 0.44 (SE ± 0.07) for BVDV. However, the heritability estimates transformed to the underlying scale (h^2^) (as calculated from Dempster and Lerner 1950) were moderate to high from 0.14 for Q-fever to 0.93 for *Brucella abortus*.

### Mapping SNP makers associated to serological response traits

Our genome-wide association study (Figure 3) identified a total of 53 SNP markers with p-values above the suggestive significant threshold (*P* > 1.49^―6^), and three of those SNP markers crossing the genome-wide significant threshold (*P* > 7.47^―8^) across all serological responses to infectious pathogens (Table 2). Thirty SNP markers mapped to several annotated genes in the *Bos taurus* genome (ARS-UCD1.2).

**Fig 3.**
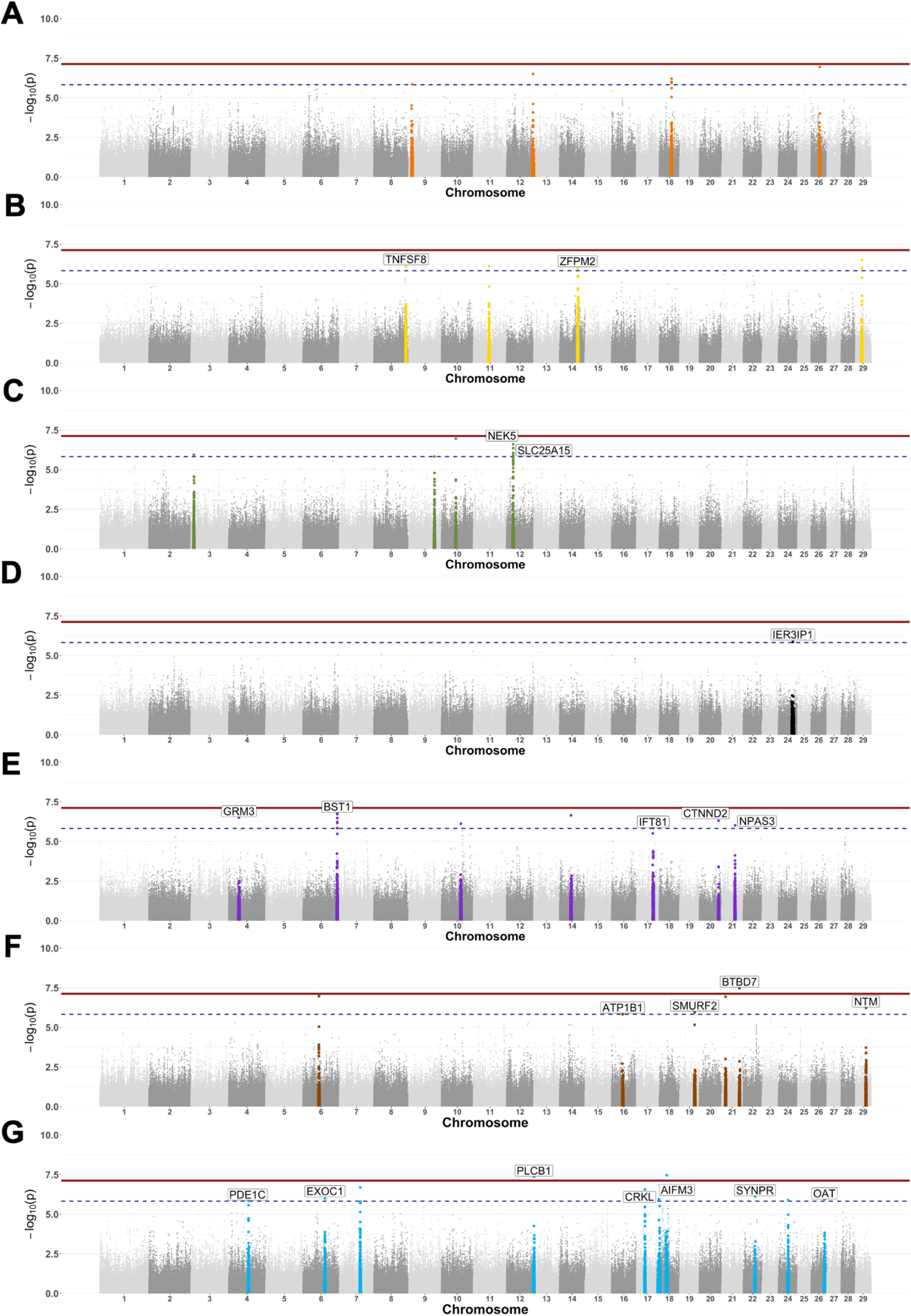
Genome-wide associations for serological response across seven infectious diseases in Tanzanian cattle. Manhattan plots show the SNP markers with the log-10 p-values above a genome-wide (7.13) and a suggestive (5.83) threshold. The adjacent 500 SNPs to those significant ones were coloured-coded to represent each of the six reproductive/zoonotic diseases: BVDV (**A**), *Neospora caninum* (**B**), *Leptospira* hardjo (**C**), RVFV (**D**), *Toxoplasma gondii* (**E**), *Coxiella burnetii* (**F**) and *Brucella abortus* (**G**).

Seven SNP makers were associated to BVDV and mapped to unannotated regions in chromosome 9,12, 18 and 26 (Fig 3A). Five SNP markers were associated to *Neospora caninum* serological response, one located within gene TNFSF8 in chromosome 8, a second SNP within gene ZFPM2, and three SNPs were located within unannotated regions in chromosome 14 and 29 (Fig 3B). Seven SNP markers were associated to *Leptospira* hardjo serological response with three SNPs located within gene NEK5 and one within gene SLC25A15. Three SNPs were located within unannotated regions in chromosome 3, 9 and 10 (Fig 3C). One SNP marker was associated to RVFV serological response which was located within gene IER3IP1 (Fig 3D). Twelve SNP markers were associated to *Toxoplasma gondii* serological response with most SNPs located genes GRM3 (one SNP), BST1 (six SNPs), IFT81 (one SNP), CTNND2 (one SNP) and NPAS3 (one SNP) located in chromosome 4, 6, 17, 20 and 21, respectively (Fig 3E). One of the SNP markers strongly associated (*P* = 3.41^e-08^) with *Coxiella burnetii* serological response was located within gene BTBD7 in chromosome 21. Three SNP associated to *Coxiella burnetii* serological response mapped within genes ATP1B1, SMURF2 and NTM in chromosomes 16, 19 and 29, respectively (Fig 3F). Fifteen SNP markers were associated to *Brucella abortus* serological response with one strongly significant (*P* = 4.30^e-08^) SNP mapping to gene PLCB1 in chromosome 13, and another SNP (*P* = 3.42^e-08^) to an unannotated region in chromosome 18. One SNP mapped within gene PDE1C in chromosome 4, three SNPs within gene EXOC1 in chromosome 6, one SNP mapped within gene CRKL and one within gene AIFM3 in chromosome 17, one SNP located within gene SYNPR in chromosome 22 and one SNP mapped within gene OAT in chromosome 26. The rest of the SNPs mapped within unannotated regions (Fig 3G). Summary statistics for SNP markers with associations at suggestive and genome-wide threshold are described in S1 Table. QQ plots for each of the seven GWAS is presented in supplementary material (S4 – S10 Figs).

## Discussion

In this study we present several important findings obtained from hard to measure phenotypes collected from smallholder dairy cattle herds across the six regions in Tanzania, mainly admixed with a high proportion of European taurine ancestry. Heritability estimates for antibody response varied with pathogen assayed but were high in BVDV, *Leptospira hardjo* and *Neospora caninum* exposure. Several SNP markers with significant association to the studied traits were identified by genome-wide association analysis. Some of the identified loci mapped to *Bos taurus* genome regions, including annotated genes involved in cell exocytosis pathways and immune response. Our GWAS results will add to the knowledge of genomic regions with potential to improve cattle resistance to infectious diseases.

The history of domestication of African cattle breeds is complex and marked by multiple admixture events between local African taurine, indicine zebu, and more recently, European (e.g., Jersey, Ayrshire and Friesian) breeds which has resulted in highly diverse populations [^34^]. As shown in our population structure and admixture analysis (Fig 2), our Tanzanian smallholder dairy cattle were crosses of European and indicine breeds, comparable to other East African smallholder cattle populations [^35,36^], but with higher levels of European introgression. The percentage of European and African taurine, and indicine zebu ancestry obtained in admixture analysis of animals was not significant when including them in both heritability and GWAS models, likely due to the few indigenous (or closely indigenous) sampled animals. As such, and given sampling ‘pure’ breeds under similar conditions is challenging, we were not able to investigate genetic differences between breeds and their effect on antibody responses which has been previously described as a key factor in bovine trypanosomiasis, tuberculosis, and East Coast fever [^32,34,37^]. In this admixed Tanzanian cattle population, however, we were able to identify several regions in the available *Bos taurus* genome with strong association to antibody response to different infectious diseases.

The prevalence of the infectious pathogens observed in this study population was variable with the highest percentage in BVDV, *N. caninum*, *Leptospira hardjo* and RVFV. These findings partially correspond to the most attributable livestock abortion agents (e.g., RVFV, *N. caninum,* and pestiviruses) found in aetiological surveys in Tanzania [^9,38^]. Importantly, our heritability estimates in the observed scale (0/1) are less accurate with the lower prevalence diseases (e.g., 2%) given the amount of variance introduced by measurement error which is reduced by transforming it to the underlying scale [^39–41^].

We identified 53 SNPs with strong association to serological response across pathogens – some which mapped to annotated genes in the *Bos taurus* genome (Table 2). The seven SNPs associated to bovine viral diarrhoea virus serological response mapped to unannotated regions (Fig 3A). Bovine viral diarrhoea virus is a highly contagious pathogen with a complex epidemiology that affects dairy cattle herds worldwide by causing persistent infection, poor reproductive performance, and huge economic loss [^42,43^]. Several BVDV control options exist (e.g., vaccination, biosecurity, removal of persistent infected animals, etc [^44^] but rarely implemented and/or maintain in LMICs [^45,46^]. With the advances of genome editing however, it has been possible to breed the first calf with reduced BVDV susceptibility by altering the CD46 gene. BVDV binds to two peptide domains in CD46 to infect cells, and therefore, an altered CD46 molecule seems to limit viral load in the blood (viraemia) in a edited calf. [^21,22^]. In the RVFV GWAS, one SNP above the suggestive significant threshold mapped within gene IER3IP1 which mutations cause a neurodevelopmental disorder in humans, and it has recently been demonstrated to play a fundamental role in B cells development in mice [^47^].

The apicomplexan parasites, *Toxoplasma gondii* and *Neospora caninum*, we studied are known for causing reproductive problems in cattle, and/or represent a high zoonotic potential (*T. gondii*) [^48^]. In our *T. gondii* serological response GWAS results, we found that ten of the twelve associated SNPs were located across annotated genes, GRM3, BST1, IFT81, CTNND2 and NPAS3 (Table 1). The GRM3 gene encodes proteins that regulate neurotransmitters (e.g., glutamate) with gene mutations directly associated to neurological disorders such as schizophrenia [^49^]. In addition, GRM3 has also been shown to suppress colon cancer and glioblastoma growth [^50,51^]. The BST1 gene encodes a molecule that facilitates pre-B-cell growth and it has been involved in autoimmune diseases and neurological disorders in humans [^52,53^]. The CTNND2 gene has been previously identified in GWAS in Nelore cattle where function may be important for growth, meat quality and milk production [^54^]. The rest of the annotated genes are involved in ciliogenesis/spermatogenesis (IFT8) and neurogenesis/schizophrenia disorder (NPAS3) [^55–57^]. Investigations of the relationship between toxoplasmosis and genes involved in neurological processes have only recently become available [^58^]. Several studies have indicated that *T. gondii* infection increases the risk of neurological disorders such as schizophrenia [^59–62^]. This risk may be somewhat explained by the potential of *T. gondii* to deregulate neurotransmitters including glutamate, which is synthetised by astrocytes that are heavily affected during *T. gondii* infection [^63^]. Therefore, it is not entirely unlikely that several of these variants within GRM3, BST1 and NPAS3 genes may be associated to host *T. gondii* infection response as revealed by our GWAS results.

**Table 1.**
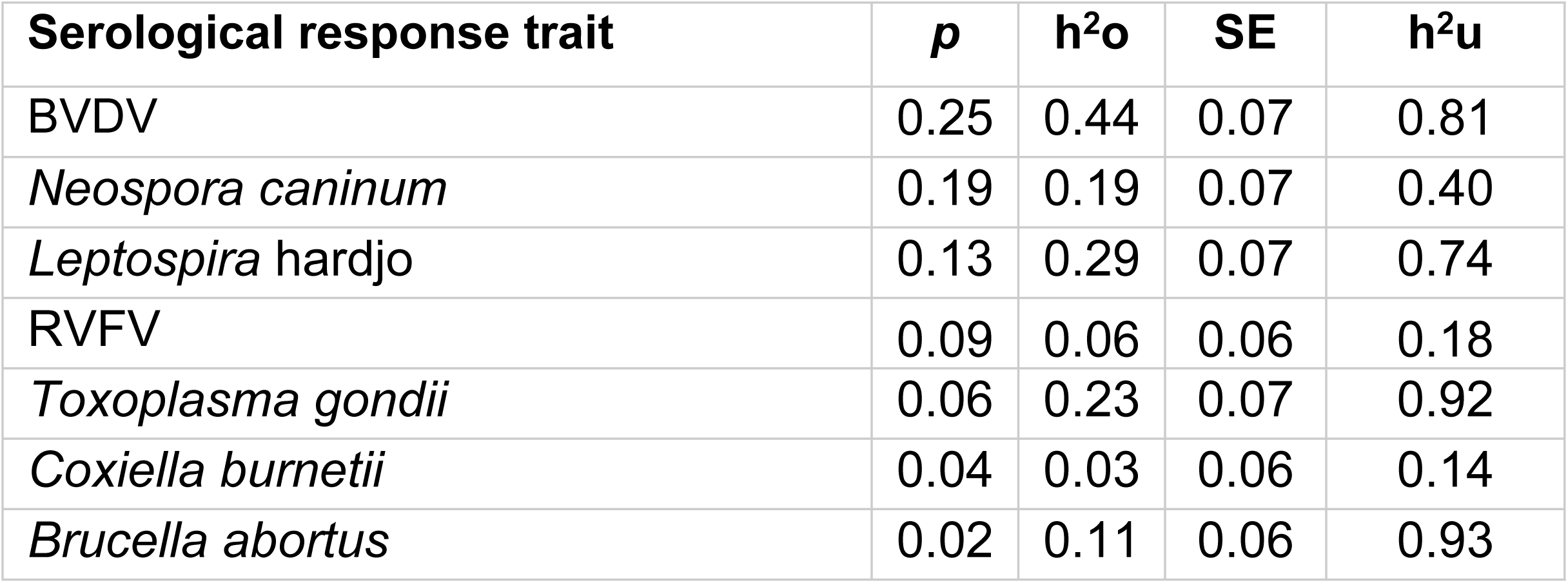
Estimated heritability for serological response to six important reproductive and zoonotic infectious diseases among Tanzanian smallholder dairy cattle. For all serological response traits, the seroprevalence (*p*), heritability on the observed scale (h^2^u) with standard errors (SE), and heritability on the underlaying scale (h^2^u) is provided.

**Table 2.**
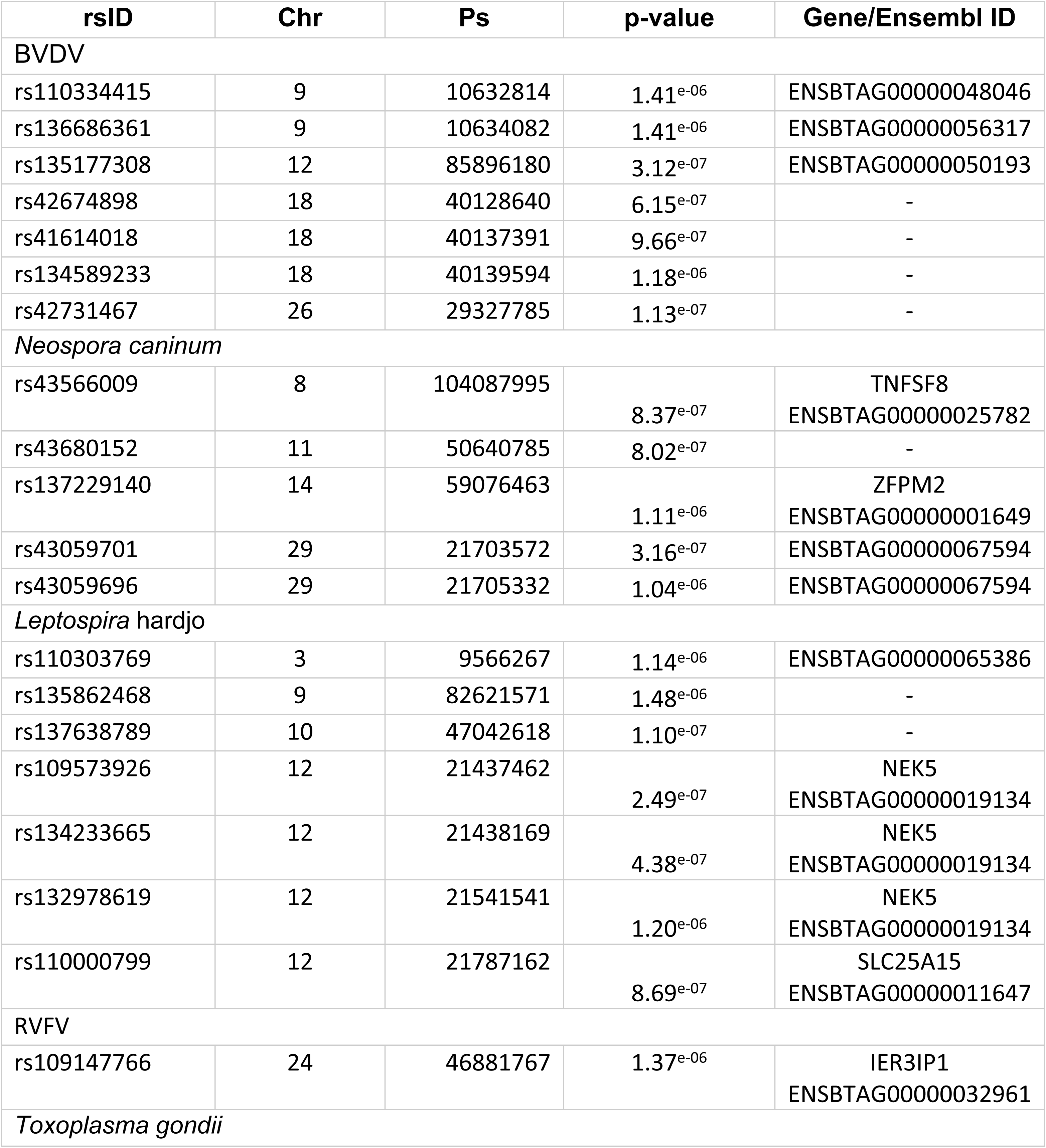
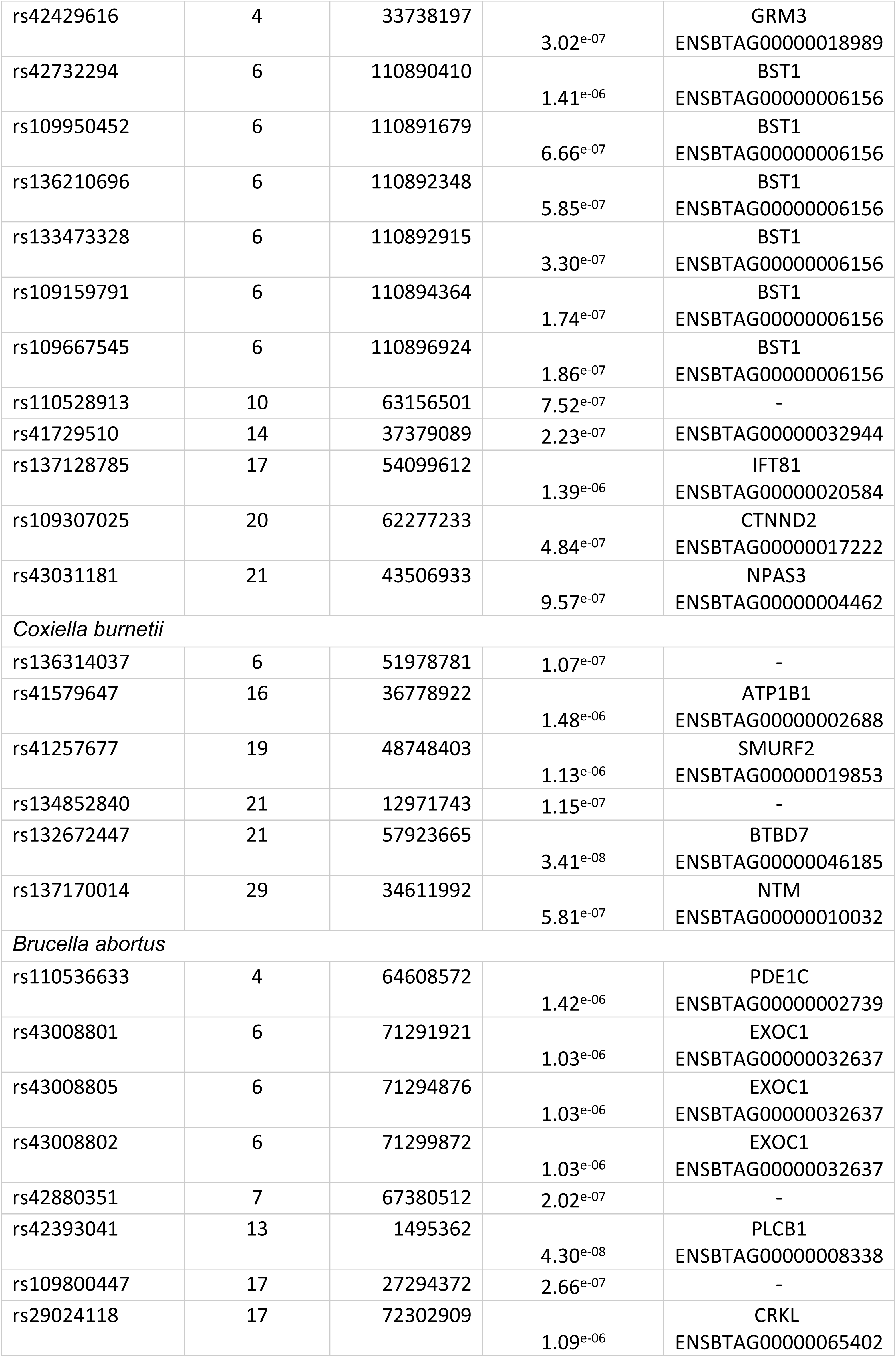
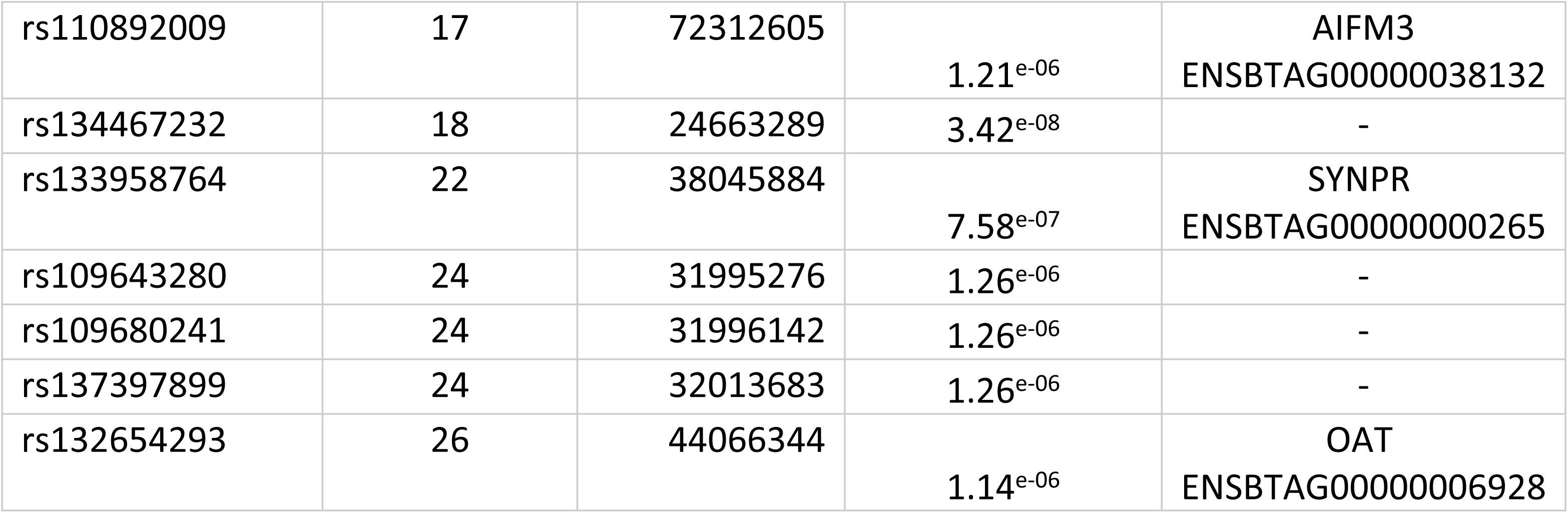
Significant SNP markers identified using linear mixed effects models in GEMMA software. For each GWAS, reference SNP cluster ID (**rsID**), chromosome (**Chr**) base-pair location (**Ps**), p-value and whether that SNP mapped to a given annotated gene (**Gene**) or Ensembl ID are provided.

In the case of *N. caninum,* we identified five SNPs – one mapped within the TNFSF8 gene, a second within the ZFPM2 gene and three SNPs were located within unannotated regions (Fig 3B). The tumour necrosis factor superfamily (TNFSF) genes encode proteins and molecules responsible for host immune defence and tumour suppression [^64,65^]. The TNFSF8 gene expression plays an important role on immune cells (e.g., CD4 T cells) defence against *Mycobacterium tuberculosis* and Hepatitis C virus infection [^66,67^]. The zinc finger protein (ZFPM2) is responsible for encoding GATA transcription factors (zinc finger DNA binding proteins) which control development of erythrocytes and immune cells such as CD4 T cells [^68,69^]. Although the link between GATA transcription factors and *N. caninum* immune response is unknown, they seemed to have an important role to local immune response against *Pseudomona aeruginosa* [^70^].

Several of the SNPs with significant association to bacterial pathogens (*L. interrogans* serovar Hardjo, *C. burnetii,* and *B. abortus*) mapped within genes NEK5, SLC25A15, ATP1B1, SMURF2, BTBD7, NMT, PDE1C, EXOC1, PLCB1, CRKL, AIFM3, SYNPR or OAT. In this *B. abortus* GWAS results, three SNPs were located within the EXOC1 gene, which is part of a complex of proteins that regulate cell exocytosis pathways. The EXOC1 encoded protein is part of the 8-molecule complex that binds the cell plasma membrane to endosomal compartments. Several bacterial organisms including *Listeria monocytogenes, Staphylococcus aureus* and *B. abortus* use different exocytic pathways to infect host cells [^71,72^]. After *B. abortus* is phagocyted by mainly macrophages or dendritic cells, it initially interacts with early endosomes to enter the cell, and subsequently replicate within the endoplasmic reticulum [^73^]. Therefore, genetic variation at the EXOC1 region may determine whether *B. abortus* infection will occur or not in a host cells. Although *C. burnetii* serological response association with NMT gene is unclear, NTM gene has been identified in association to *B. abortus* infection in wild boars, and displaced abomasum disorder in Holstein cattle [^74,75^]. The association between SNPs located within BTBD7 and PLCB1 genes and serological response against bacterial pathogens is not clear, however, BTBD7 gene has been associated to indicators of heat stress in Holstein cattle as well as involved in other biological processes such as development and tumour progression [^76,77^]. The NEK5 and SLC25A15 genes encode proteins involve in mitochondrial functions, cell-cycle progression and metabolism, and tumorigenesis, but without a clear role in immune response to bacterial infection [^78,79^]. In vitro human gene expression experiments have showed that ATPase Na+/K+ transporting subunit β 1 (ATP1B1) protein limits DNA and RNA viruses’ expression and replication by promoting IFNs and proinflammatory cytokines activation [^80^]. Although the SMAD specific E3 ubiquitin protein ligase 2 (SMURF2) has a primary role in TGF-β signaling pathways (e.g. embryogenesis, cellular homeostasis, etc), it has recently been shown to affect anti-viral signaling pathways (e.g., Type 1 IFN signaling, binding filovirus VP40 matrix proteins) [^81,82^]. The PDE1C, CRKL, AIFM3, SYNPR and OAT genes are involved in developmental and metabolic functions; however, their role in immune response to bacterial pathogens has not yet been studied.

While our study provides genetic estimates and associations of antibody response to several pathogens among admixed African cattle populations, we realise our sample size is a limiting factor for our GWAS analyses. However, it should be noted that collecting this kind of phenotypes in smallholder dairy units is labour intensive and expensive. Our data identified regions in the bovine genome involved in host immune defence and pathogen infection which warrant further investigation e.g. in vitro experiments of gene expression or genome editing as in the case of the BVDV edited calf.

## Materials and Methods

### Sample collection and study area

Genotypes for the cattle were obtained from the African Dairy Genetics Gains (ADGG) project (https://www.ilri.org/research/projects/african-dairy-genetic-gains) which consisted of 2045 crossbred Tanzanian cows and bulls sampled from their larger database. ADGG has managed a performance (e.g., milk yield, body weight, etc) and genotype database for most of the registered animals, mainly smallholder cattle, from Tanzania and other low-and middle-income countries (LMICs) since 2016. This cattle subset was selected based on confirmed presence in both the household and the ADGG database at the time of collection. Households were located across twenty-three districts in six regions of dairy production importance in Tanzania [^1,2^] (Fig 1).

A cross-sectional survey and sample collection was carried out by our veterinary team, ADGG and Tanzanian Livestock Research Institute (TALIRI) staff from July 2019 to October 2020. Information on households (e.g., geographic location), animals (e.g., age, sex, phenotypical features, etc), and herd management (e.g., feeding, reproduction, and health) were recorded electronically using the open data kit (ODK) platform, and curated for downstream analysis using R and RStudio [^83,84^].

Blood samples were collected from each animal by jugular venopuncture using plain vacutainer tubes (BD vacutainer®, Auckland, New Zealand), centrifuged and refrigerated until processing at regional laboratories across the study regions in Tanzania. Serum was aliquoted in cryovial tubes and stored at -20 °C at the Nelson Mandela African Institution of Science and Technology (NM-AIST) in Arusha, Tanzania.

Ethics of the study for animal subjects was reviewed and approved by the International Livestock Research Institute Institutional Animal Care and Use Committee (ILRI-IACUC2018-27) and the research permit was granted by the Tanzania Commission for Science and Technology (COSTECH), Ref. (2019-207-NA-2019-95).

### Description of Tanzanian cattle population

On average cattle in this study were five years of age with 97.2% being females. Phenotypic characterisation classified 3.8% as having African indigenous features, whereas 96.2% were identified as crosses between East African Shorthorn Zebu and Ayrshire, Holstein or Jersey breeds. Management strategies in this population varied with most animals placed under an intensive feeding system with little opportunity to graze freely in open pastures. Reproductive management was carried out through artificial insemination, and a small percentage of farmers reported the use of an owned or hired bull as a mode of reproduction in the herd. Preventative disease control measures were carried out through vaccination, although it was not routinely implemented with only 15.2% of farmers reporting using mainly a foot-and-mouth disease vaccine. A total of 23 Districts were sampled with the number of animals in each district varying from 15 in Iringa Municipal Council to 261 in Moshi Rural District Council. The herd size, composed mainly of heifers or cows, was variable with only a few herds larger than 50 mature females.

### Serological health resilience traits

The serological response trait was obtained for each individual animal through testing for antibody response to seven pathogens using commercial ELISA kits of bovine viral diarrhoea virus (ID Screen® BVD p80 Antibody Competition, Innovate Diagnostics, France), *Neospora caninum* (ID Screen® *Neospora caninum* Competition, Innovate Diagnostics, France), *Leptospira hardjo* (*Leptospira interrogans* subtype Hardjoprajitno and *Leptospira borgpetersenii* subtype Hardjobovis; The Linnodee Leptospira Hardjo ELISA Kit^TM^, Linnodee Animal Care, UK), Rift Valley fever virus (ID Screen® Rift Valley Fever Competition Multi-species, Innovate Diagnostics, France), *Toxoplasma gondii* (ID Screen® Toxoplasmosis Indirect Multi-species, Innovate Diagnostics, France), *Coxiella burnetii* (Q fever; PrioCHECKIT™ Ruminant Q Fever Ab Plate Kit, Thermofisher Scientific, USA) and *Brucella abortus* (COMPELISA 160 & 400, APHA Scientific, UK). Following manufacturer guidelines (see details in [^6–8,85^]. All OD values were transformed into a binary seropositive seronegative classification based on the manufacturers recommended cut-offs.

### Genotyping and imputation

Following a method previously applied to East African crossbreed dairy cattle [^35^], we genotyped our samples using a low-density GeneSeek Genomic Profiler Bovine 100K chip and inferred missing genotypes using a reference cattle population genotyped with a high-density chip. As described in [^35^], the imputation method uses a crossbreed cattle population, mainly from East Africa (see details in [^35^] Table 1), and a ‘pure’ breed reference (British Fresian, Holstein, Jersey, Guernsey, Nelore, and N’Dama) population (n = 3091), both genotyped using the Illumina BovineHD BeadChip (Illuimina, San Diego, CA). This reference population was provided by authors in [^35^] and was solely use for imputation. In this procedure, autosomal SNPs with a GC score > 15%, a call rate > 90% and a minor allele frequency (MAF) > 0.01, and animals with below 10% missing genotypes were kept for the imputation pipeline. Pre-phasing and imputation of genotypes were carried out using Eagle v2.4.1 [^86^] and Minimac3 v2.0.1 [^87^] programmes, respectively.

### Genomic differentiation and ancestry estimation

To explore population structure and ancestry in our samples, we first merged our samples to a reference population [^32^]. We then ran principal component analysis (PCA) and model-based estimation of ancestry on the merged genotype data. Our reference population included closely pure European (ET, n=99; 63 Holstein and 36 Jersey) and African taurine (AT, n= 59; 47 N’Dama and 12 Muturu), and Asian zebu (AZ, n= 65; 30 Gir and 25 Nelore) cattle genotyped with the Illumina BovineHD BeadChip [^32^]. Both datasets were converted from TOP to FOR format and aligned using the SNPchiM v3 programme [^88^], and subsequently merged as a single VCF file using the bcftools v1.3 suite [^89^]. The merged data was converted from VCF to BED format using the Plink v1.90 programme [^90,91^] in which SNPs with a MAF < 0.01 (--maf) and a call rate > 90% (--geno), as well as animals with above 10% missing calls (--mind) were removed. Additionally, we removed animals with a degree of relatedness above 0.25 (parent-offspring and full siblings) based on KING-robust estimator [^92^] which has been implemented in the Plink v2.0 programme [^90,91^].

Population structure between our Tanzanian and reference samples was explored by performing a principal component analysis (PCA) on the genotype data using Plink v1.90. We also ran a separate PCA with only the Tanzanian samples with the first five PCs later being used as covariates in models for genetic parameter estimates and genome-wide association analysis. Prior to ancestry estimation, SNP markers in high linkage disequilibrium (LD) were removed after applying a r-squared threshold > 0.2 with another SNP within a 200-SNP window with sliding windows of 10 SNPs at a time. To estimate the level of ET, AT and AZ ancestry in our Tanzanian cattle, we performed a supervised ADMIXTURE [^93^] analysis using a 5-step Expectation-Maximization (EM) algorithm. We ran a 10-fold cross-validation with 200 bootstrap resampling to calculate standard errors assuming *K* = 2 to *K* = 3 fixed ancestries. We compared our supervised analysis with an unsupervised ADMIXTURE analysis with the same parameter setting but exploring *K* = 2 to *K* = 23 clusters (S2 Fig). Q matrix estimates for each cluster were visualised using the R *pophelper* package [^94^] and the best value of *K* was chosen based on the lowest cross-validation error in unsupervised analysis.

### Genetic parameter estimation

An univariate linear mixed effects models (LMM) was fitted in each of the six serological traits and their heritability were obtained using Restricted Maximum Likelihood (REML) implemented with the ASReml software [^95^]. The fitted LMM included fixed effects of sex (male and female), sample collection month (seven months), districts (23 districts), herd size (four categories) and the first five PCs calculated with the genotype data as covariates. An animal random effect was fitted using a genomic relationship matrix (GRM) computed with the genotype information using method 2 of VanRaden *et al* [^96^]. The following model was fitted:

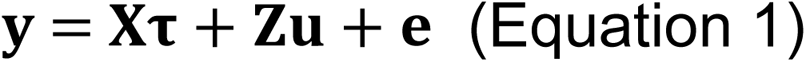

Where *y* is the vector of observations (serological traits; that is sero-response to BVDV, *N. caninum*, *L. interrogans* serovar Hardjo, RVFV, *T. gondii*, *C. burnetii* or *B. abortus*) modelled on the observed scale (0-1) assuming that data as continuous, **τ** is a vector of fixed effects, *u* is a vector with the additive genetic effects, *X* and *Z* are design matrices associating observations to fixed and random effects, respectively, and *e* is the vector of residual errors.

The narrow-sense heritability (ℎ_0,1_^2^) on the observed scale is estimated as the proportion of phenotypic variance (*σ_P_*^2^) explained by the additive genetic variance (*σ_P_*^2^).

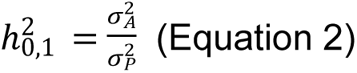

We then calculated the under the ‘underlying’ (ℎ*_u_*^2^) scale based on Dempster & Lerner [^40,41^].

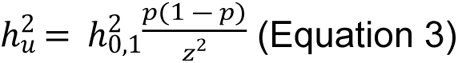

Where *p* is the population prevalence and *z*^2^ is the ordinate of the standardized normal curve corresponding to a probability *p*. In practice, a binary disease trait (0/1) is defined by exceeding or failing a threshold (heritability on the observed scale) and the heritability in the underlying scale take into account that definition depends on the disease prevalence [^39,40^].

### Genome-wide association analysis

A genome-wide association study (GWAS) was carried out on the six traits separately with the GEMMA software [^97^] fitting LMM with the same effects included model used in the variance component analysis described above (animal sex, sample collection month, districts, herd size as fixed effects, five PCs as covariate and animal genetic effect as random)plus the genotype effect of the SNP being tested animal (GRM). To account for multiple testing, the significant threshold for the SNP effects were adjusted using Bonferroni correction, the SNP effects were declared significant at the suggestive of genome-wide levels when their log-10 p-value was greater than 5.83 and 7.13, respectively. SNP log-10 p-values were visualised in Manhattan (Fig 3) and QQ (S4 – S10 Figs) plots.

To further calculate allele substitution effects, SNPs which crossed genome-wide significant thresholds and suggestive thresholds were included as fixed effects, together with the previously fitted fixed effects in ASReml to estimate their additive and dominance effects. The genotypic means were defined as AA, AB and BB from the predicted trait value for each genotype class, *p* and *q* as the SNP allele frequencies, and VA as the total trait additive genetic variance when no SNP effects are included in the model. The genetic effects were then calculated as follow: additive effect, *a* = (AA – BB)/2; dominance effect, *d* = AB – [(AA + BB)/2]; and proportion of genetic variance due to SNP = [2*pq*(*a*+*d*(*q* -– *p*))^2^]/VA.

### Genome mapping of associated loci

In order to find the genes that may be responsible to serological response in our Tanzania cattle we searched SNP makers found in GWAS in the annotated *Bos taurus* ARS-UCD1.2 genome in Ensembl.org.

## Acknowledgements

We would like to acknowledge David Wragg for help and support on merging reference and this study cattle genotypes, and Hassan Aliloo for his guidance on genotype imputation.

## Author contributions

E.A.J.C., B.M.C.B., and G.M.S. designed the study.

L.E.H.C. and B.M.C.B. wrote the manuscript with contributions from E.A.J.C., O.M., and G.M.S. I.M., S.K.M., S.B., B.Z.S., D.W., R.P.W., J.P., R.M., P.G.T., D.M.K., E.L., and O.A.M. revised and edited the manuscript.

I.M., S.K.M., S.B., D.M.K., and E.L. collected cattle samples in Tanzania.

I.M., S.K.M., and S.B., B.Z.S., B.K., G.N., and E.A.J.C. carried out serological laboratory activities.

R.M., and O.A.M. curated and provided cattle Illumina genotyping data for this project.

L.E.H.C analysed the data with contributions from H.A SNP array imputation, D.W. in merging sample and reference cattle genotypes, R.P.W. in estimating a genomic relationship matrix, and O.M. guiding genome-wide association analysis.

## Supporting information

### Data availability

Raw sequenced data will be uploaded to the repository on publication.

### Materials & Correspondance

Correspondence to Mark Bronsvoort and Luis Enrique Hernandez Castro

### Code availability

Code for admixture and GWAS analyses will be available via Github repository (github.com/lehernandezc/tanzania_gwas) on publication.

